# Running against the clock: describing microbial diversity in an extremely endangered microbial oasis in the Chihuahuan desert

**DOI:** 10.1101/2024.09.04.611308

**Authors:** Ulises E. Rodriguez-Cruz, Manuel Ochoa-Sánchez, Luis E. Eguiarte, Valeria Souza

**Affiliations:** Departamento de Ecología Evolutiva, Instituto de Ecología, Universidad Nacional Autónoma de México, Ciudad de México, Mexico; Doctorado en Ciencias Biomédicas, Universidad Nacional Autónoma de México, Ciudad de México, México; Doctorado en Ciencias Biológicas, Universidad Nacional Autónoma de México, Ciudad de México, México; Centro de Estudios del Cuaternario de Fuego-Patagonia y Antártica (CEQUA), Punta Arenas, Chile

## Abstract

Cuatro cienegas Basin (CCB) is an extraordinary diverse oasis that is dying in front of our eyes. The Churince system disappeared completely in 2016 and the poza named for its amazing diversity, Archaean Domes, disappeared in 2023. However, there could be groundwater water connections between sites, allowing to dispersal of taxa as well as the survival of microbial lineages in the deep aquifer Herein, we explore whether there is microbial connectivity across sites in the oasis of CCB, located in the Chihuahuan Desert, in the Northern state of Coahuila in Mexico. We focused first on two distinct aquatic system within CCB that are ∼4700 meters apart on the Eastern lobe of the basin: Pozas Rojas –a preserved system characterized by several unique ponds surrounding a deep lagoon, and the now extinct Archean Domes (AD)--noted for its extreme salinity, as well as anoxic and methane-rich conditions. We sequenced the hypervariable V4 region of the 16S rRNA gene from samples of Pozas Rojas (PR), and of two different features in AD: orange circles (C) and dome-shaped structures (D). Additionally, we utilized whole-genome metagenomic data from different samples from AD site (from the years 2016 to 2023). Second, we compared metagenomes obtained from the now extinct Churince (samples from 2012 to 2014) in the West lobe of the basin, with the Easter side. Our results revealed taxa common to different sites, including *Halanaerobium* sp. and *Desulfovermiculus* sp., that are prevalent across samples from PR and AD, and shared between AD and Churince, suggesting that those genera are core to the deep aquifer.

Moreover, during the eight years of sampling, AD exhibited a substantial decline in microbial diversity. Possibly, this decline, as well as Churince demise, is due to anthropogenic disturbances of the deep aquifer due to overexploitation for alfalfa irrigation. Diversity in metagenome-assembled genomes (MAGs) decreasing from 125 MAGs in 2016 to only 22 in 2023. Our study supports the hypothesis of the importance of groundwater as a fundamental microbial reservoir and connector among CCB ponds. Therefore, the regulation of extraction of such aquifer is paramount.

## 1.0 Introduction

Considering that CCB is an endangered complex system of ponds connected by underground water(Wolaver et al., 2013) that harbors a “lost world” of unique microbial diversity (Souza et al., 2018). It is possible that such ancient lineages have survived eons inside the deep aquifer of the “San Marcos y Pinos Sierra”. We believe that the demise of the superficial microbial diversity does not imply extinction *per se*, but a lack of understanding of the deep aquifer dynamics in relation to the surface ponds. So far, the beta diversity among sampling points even at a small scale of cm (Espinosa-Asuar et al., 2022) is very high, suggesting that there is a subsampling of a much bigger seed bank rooted in the deep aquifer. Therefore, despite been overexploited, such deep aquifer may hide and protect CCB ancient lineages from extinction. In which case, the oasis microbial diversity can be recovered with the proper policy toward the exploitation of its water recourses.

We already know that the composition of microbial communities inhabiting water bodies is driven by seasonality in lakes and marine surface ecosystems (Shade et al., 2012;Ward et al., 2017; Wang et al., 2020). However, marine microbiome composition below the euphotic zone shows a stable microbial pattern despite seasonality, highlighting the role of solar light and temperature in the structure of aquatic microbiomes (Ibarbalz et al., 2019). Hence, in water ecosystems devoid of light and solar radiation, such as groundwater, it is expected that microbial communities might be stable through time but different among sites due to superficial environmental heterogeneity (GRIEBLER & LUEDERS, 2009) .Groundwater harbors a vast portion of microbial diversity, which is influenced by features of the groundwater ecosystem (e.g., rock type), microbial interactions, biogeochemical cycles, and human perturbation (Fillinger et al., 2023) .Interestingly, microbial dispersion could occur between groundwater and surface ponds (Ji et al., 2022). Yet, microbial dispersion is hindered by surface environmental stress, although this limitation varies across phylogenetic groups which means that some bacteria are more vulnerable to being locally restricted due to stress than others (Gruzdev et al., 2023; Ning et al., 2024). However, whether microbial dispersion occurs among freshwater ponds with varying oligotrophic conditions but sharing a deep aquifer, has not been explored.

Several aquatic systems in CCB have been thoroughly researched (Souza et al, 2018). Among them, stands out the Archean domes (see Fig 1) (AD, hereafter) (see Espinosa-Asuar et al., 2022; Medina-Chávez et al., 2023; Madrigal-Trejo et al., 2023). In this site, A pH of 9.8 and a salinity of 5.28% were measured during the rainy season, while in the dry season, the pH decreased to 5 and the salinity increased to reach saturation. During the rainy season, a stoichiometric imbalance of C:N:P of 122:42:1 was reported (Espinosa-Asuar et al., 2022; Medina-Chávez et al., 2019; Madrigal-Trejo et al., 2023; Rodriguez-Cruz, et al., 2024). AD microbial diversity undergoes taxonomic changes across seasons, yet its functionality remains similar. Interestingly, most of the community was shared across sampled seasons, highlighting the presence of core taxa, despite environmental fluctuations (Madrigal-Trejo et al., 2023). Similarly, viral communities do not appear to be influenced by environmental fluctuations; rather, their composition remains similar across seasons and years (Cisneros-Martínez et al., 2023). Interestingly, AD had a high diversity of Archaea (Medina-Chavez et al., 2023; Rodriguez-Cruz et al. 2024). In what we believe to be a deep aquifer surgence in 2019, Archaea prevalence in the surface moved from ∼5% numbers to above ∼30% (Madrigal Trejo et al., 2023). Most of these Archaea are endemic to AD and represent phylogenetically basal lineages (Rodriguez-Cruz et al. 2024).

**Figure 1.**
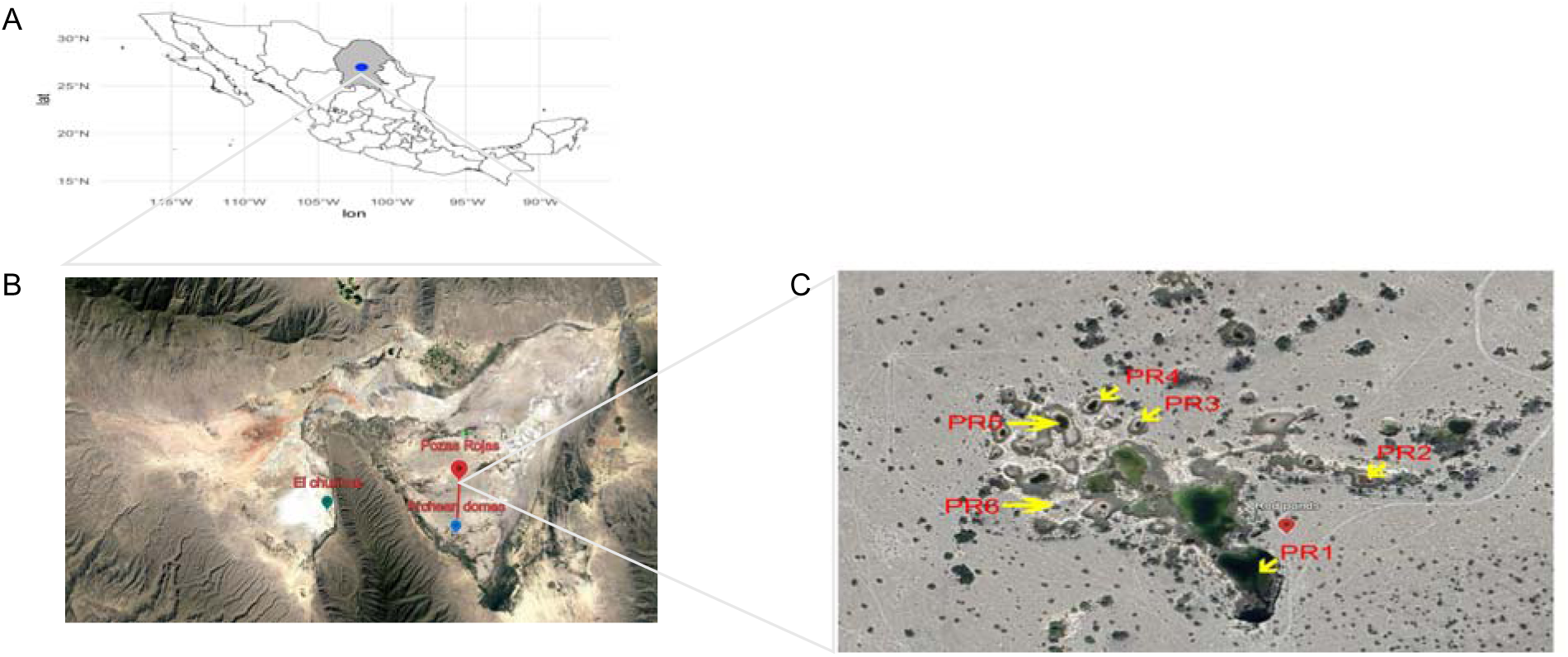
(A) Map showing Coahuila state in northern Mexico, with a blue point indicating the location of the Cuatro Ciénegas Basin (CCB). (B) Aerial view of the CCB. (C) Distribution of the Pozas Rojas ponds within the CCB**. Map data: ©2024 Google, TerraMetrics**

On the other hand, the Churince flow system is in the western region of the CCB. It was thoroughly studied by us from 2000 to 2016 when it totally dried out (Souza et al., 2018). It was particularly interesting system given its strong gradient for salinity, temperature, pH and dissolved oxygen (Johannesson et al. 2004; Carson and Dowling 2006). Besides, Churince was unique within CCB, since it is surrounded by large and pure gypsum dunes (Szynkiewicz et al. 2010). This site consisted of a freshwater spring that connects to an intermediate shallow pond via a small stream and eventually terminated in a very large shallow lagoon that was part of the dynamic of the gypsum dunes. The environment was extremely poor in phosphorus (concentrations of PO^-4^ lower than 0.1 μmol (Elser et al. 2005) but rich in sulfate and magnesium. However, starting in late 2002, the pumping of its subterranean water by nearby alfalfa ranchers completely drained the large lagoon by 2008 and the intermediate lagoon by 2016.

We have hope that Pozas Rojas, the only well-preserved system in CCB, can work as a kind of “lifesaver”. This is a site where the dynamics between the deep aquifer and surface water are still maintained and microbes that depend on light can survive as well as the deep biosphere.

On one hand, we hypothesized that natural water circulation between the deep aquifer and the surface water in the Cuatro Ciénegas Basin (CCB), as described by Wolaver et al. (2012), promotes the dispersal of microbiological diversity across the region. As such, we predict the presence of core microbial taxa shared among different ponds, despite their different surface environmental conditions. To test this, we sequenced the V4 hypervariable region of the 16S rRNA gene from samples collected at two sites: Archean Domes (AD) and Pozas Rojas (PR), on the East side of the basin, separated by 4700 meters. Our results revealed distinct spatial patterns in microbial community composition, suggesting potential connectivity between the ponds.

On the other hand, to test the hypothesis of microbial dispersion at a larger scale throughout the Cuatro Ciénegas Basin (CCB), we analyzed whole-genome metagenomic data from the AD site collected between 2016 and 2022 (Rodríguez-Cruz et al., 2024) alongside unpublished data from the same site in 2023. This dataset was compared with data from another aquatic system within CCB, Churince. The Churince data were obtained from De Anda et al. (2018), covering the period from 2012 to 2014 (dates where the system was already stressed) from which they obtained 12 metagenomes. The aim of this comparison was to identify a shared microbiome between the two sites across different detection thresholds and relative abundance levels. Additionally, we attempted to assemble metagenome-assembled genomes (MAGs) from Churince to assess temporal changes in microbial diversity. However, due to variations in MAG quality across years, a comprehensive analysis of MAG gain and loss over time was not possible.

Our main finding reveals that several core genera are prevalent across sites, with *Halanaerobium* sp. and *Desulfovermiculus* sp. being particularly significant suggesting at least some potential dispersal between sites along the deep aquifer as suggested previously by Ulloa et al., 2022 studying water microbial diversity of Pozas Rojas vs Churince.

## 2.0 Methodology

### 2.1 Sampling

From 2020 to 2023, samples were collected in AD in both the dome-forming (D) and the orange circle-forming zones (C) (26°49′41.7″N, 102°01′28.7″W) (see Fig. 1 for sampling details) at various depths, ranging from the surface to approximately 50 cm in depth. A total of 19 samples were collected: 12 samples of dome zone and seven of orange circle zone. In addition, we sampled in 2020, 2021 and 2023 six ponds in the PR system (26°52′N, 102°1′W, see additional sampling details in Fig. 1 and in Table 1). Sampling was conducted under the collection permit SGPA/DGVS/03188/20 issued by the Subsecretaría de Gestión para la protección Ambiental, Dirección General de Vida Silvestre.

Following collection, the samples were immediately transferred to liquid nitrogen and stored until total DNA extraction.

**Table 1.** Distance in kilometers between each pond in the PR aquatic system of CCB.

|  | PR1 | PR2 | PR3 | PR4 | PR5 | PR6 |
| --- | --- | --- | --- | --- | --- | --- |
| PR1 | 0 | 0.096 | 0.113 | 0.144 | 0.146 | 0.132 |
| PR2 |  | 0 | 0.126 | 0.161 | 0.178 | 0.190 |
| PR3 |  |  | 0 | 0.035 | 0.053 | 0.082 |
| PR4 |  |  |  | 0 | 0.026 | 0.072 |
| PR5 |  |  |  |  | 0 | 0.049 |
| PR6 |  |  |  |  |  | 0 |

### 2.2 Total DNA extraction and amplicons sequencing

The DNA extraction followed a column-based protocol with a Fast DNA Spin Kit for Soil (MP Biomedical). DNA was sequenced at the Laboratorio de Servicios Genómicos, LANGEBIO (http://langebio.cinvestav.mx/labsergen/) using Illumina MiSeq 2x300 bp for the V4 region of the 16S rDNA gene (515F universal primer 5′-GTGYCAGCMGCCGCGGTAA-3′/ 806R universal primer 5′-GGACTACNVGGGTWTCTAAT′ (Walters et al., 2015) For each sample, 3 µg of genomic DNA (DO 260/280 1.8) was used.

### 2.3 Sequencing Data Preprocessing

The 16S rDNA gene sequence data underwent initial quality control procedures. FastQC (v0.11.8) (Andrews, 2010) was employed to assess raw data quality. Subsequently, the cutadapt tool (v3.2) (Martin 2011), integrated directly within the R script, was used for trimming and filtering of low-quality bases. The impact of this quality trimming on the distribution of read lengths was then evaluated to ensure the suitability of the data for further analysis. Following this, the required R libraries, including DADA2 (V.1.32.0) (Callahan et al., 2016), were loaded, and the trimmed and filtered sequencing data were imported into R as a DADA2-compatible object.

Denoising procedures were carried out using the DADA2 package. We used the filterAndTrim function for additional filtering and trimming based on quality. The learnErrors function was then used to estimate and model error rates in the data. Denoising was accomplished using the dada function, allowing for the inference of exact sequence variants (Amplicon Sequence Variants, ASVs hereafter). This step aimed to improve the accuracy of the representation of biological sequences by addressing sequencing errors. Following denoising, the identification and removal of chimeric sequences were done using the removeBimeraDenovo function. This ensured that only non-chimeric sequences contributed to subsequent analyses. Subsequently, a non-chimeric sequence table was generated for taxonomic assignment and additional analyses.

Taxonomies were assigned to the resulting ASVs using the IdTaxa function within the DECIPHER(v2.0) (Wright et al., 2012)package in the R script. The SILVA_SSU_r138_2019 database reference was specified. The non-chimeric sequences and their associated taxonomies were combined into a final table directly.

### 2.5 Alpha and Beta Diversity Analysis of the 16S rRNA Sequence Data

For the 16S rRNA sequence data, the phyloseq package in R was used to assess alpha diversity with Chao1 and Shannon indices, which provide information on both the richness and diversity of microbial communities, respectively. The analysis was conducted using the plot_richness function from the phyloseq package, which allows for the visualization of diversity indices based on the specified variables. In this analysis, the object ASV_physeq, which contains the ASV data, was used as input for the function.

16S rRNA sequencing data Beta diversity analysis was performed using the DESeq2 package (v1.44) in R (Love et al., 2014). Raw sequencing reads were transformed into a count table and metadata detailing sample grouping was incorporated. A DESeqDataSet object was created, specifying the experimental design, and subsequent normalization and variance-stabilizing transformations were applied to address variations in library sizes.

The dissimilarity matrix, representing pairwise distances between samples, was calculated based on the transformed count data using Euclidean distance metrics with the hclust (Guénard & Legendre, 2022) function in R Hierarchical clustering was employed with the Ward’s method to group samples according to their beta diversity profiles.

To investigate further the beta diversity, we employed a two-step approach. Firstly, the count data was transformed to proportions using the transform_sample_counts function from the phyloseq package (v1.48.0) (McMurdie & Holmes, 2013). This step ensures that subsequent analyses are not biased by differences in sequencing depth between samples, a common practice when working with compositional microbiome data.

Secondly, non-metric multidimensional scaling (NMDS) was performed on the Bray-Curtis dissimilarity matrix, capturing the compositional dissimilarities between samples. The plot_ordination function, also from the phyloseq package, facilitated the visualization of the NMDS results. Each sample point in the ordination plot represents a unique microbial community, with distances between points reflecting the degree of dissimilarity based on Bray-Curtis distances. To enhance the interpretability of the NMDS plot, samples were color-coded based on the location of each sample. This allowed for the exploration of spatial patterns in beta diversity, revealing potential differences in microbial community composition across different locations. The resulting plot provides a visual representation of beta diversity patterns, aiding in the identification of underlying structure and relationships within the microbial communities across the sampled locations.

### 2.5 Change in relative abundance in different detection thresholds between PR and AD

To visualize the core-microbiome, we used the phyloseq package in R. First, the detection of each taxon was calculated, defined as the proportion of samples in which the relative abundance exceeded 1%. The prevalence of each taxon was then calculated, and a minimum threshold of 10% was applied to select the ASVs of interest. Then the data were normalized, by transforming them into proportions. Finally, a heatmap of the core microbiome was generated using the plot_core function, specifying prevalence thresholds (0%, 10%, 20%, 30%, 40%, 50%, 60%, 70%, 80%, 90%, 100%) and detection thresholds (0.001, 0.003, 0.009). This approach provided a clear and detailed visualization of the core microbiome, facilitating the identification of taxa and their relative prevalence in the analyzed samples.

### 2.6 Microbial communities’ comparison between Churince and AD sites using whole genome metagenome sequencing

To further assess and compare the diversity within and among sites in CCB, we used shotgun metagenomic sequencing data for two sites: AD and Churince. From these sites, we used metagenomes previously published for AD (22 metagenomes) by Rodríguez-Cruz et al. (2024) plus, three previously unpublished metagenomes obtained in 2023. For details on DNA extraction and sequencing, see Rodríguez-Cruz et al., 2024. For Churince (26°50’55.5“N 102°08’43.9”W), we used data published by De Anda et al. (2018), which consists of 12 metagenomes from water samples collected at the Churince site between 2012 and 2014. Refer to Table 2 for more information about each sample.

**Table 2.** Sample codes taken from 2016 to 2023 from the CCB AD Pond, and metadata associated to data coming from the Churince site from 2012 to 2014.

| Sample | Type | Depth<br>sampling<br>(cm) | sampling<br>zone | Sampling<br>year | Season | raw_reads | trimmed_reads | N50 |
| --- | --- | --- | --- | --- | --- | --- | --- | --- |
| D1M01 | Microbial<br>mat | 0-10 | Dome,<br>Site DA | Apr-2016 | Dry | 28 859 454 | 26 799 269 | 2073 |
| D1M02 | Microbial<br>mat | 0-10 | Dome,<br>Site DA | Oct-2016 | Wet | 4772053 | 4340057 | 1821 |
| D1M03 | Microbial<br>mat | 0-10 | Dome,<br>Site DA | Feb-2017 | Dry | 8203484 | 7269569 | 1864 |
| D1M04 | Microbial<br>mat | 0-10 | Dome,<br>Site DA | Oct-2018 | Wet | 10030782 | 8346670 | 1401 |
| D1M05 | Microbial<br>mat | 0-10 | Dome,<br>Site DA | Mar-2019 | Dry | 25873990 | 22935865 | 1726 |
| D1M06 | Microbial<br>mat | 0-10 | Dome,<br>Site DA | Sep-2019 | Wet | 20153088 | 18787825 | 1532 |
| C1M08 | Water | 0-10 | Circle<br>Site AD | Oct-2020 | Wet | 24065589 | 19275870 | 1667 |
| C4M09 | Sediment | 30-40 | Circle<br>Site AD | Oct-2020 | Wet | 14315374 | 13361648 | 1152 |
| C5M10 | Sediment | 40-50 | Circle<br>Site AD | Oct-2020 | Wet | 18050094 | 16518645 | 1138 |
| D1M07 | Microbial<br>mat | 0-10 | Dome,<br>Site DA | Oct-2020 | Wet | 17148993 | 15124218 | 1532 |
| D4M11 | Microbial<br>mat | 30-40 | Dome,<br>Site DA | Oct-2020 | Wet | 18976795 | 17298109 | 1354 |
| D5M12 | Microbial<br>mat | 40-50 | Dome,<br>Site DA | Oct-2020 | Wet | 16106607 | 14999646 | 1487 |
| D1M13 | Microbial<br>mat | 0-10 | Dome,<br>Site DA | Sep-2021 | Dry | 11669164 | 7060584 | 1480 |
| D2M14 | Microbial<br>mat | 010-20 | Dome,<br>Site DA | Sep-2021 | Dry | 6280722 | 5893600 | 1056 |
| D3M15 | Microbial<br>mat | 0-30 | Dome,<br>Site DA | Sep-2021 | Dry | 5849741 | 3642579 | 1065 |
| D4M16 | Microbial<br>mat | 30-40 | Dome,<br>Site DA | Sep-2021 | Dry | 4894276 | 4657996 | 755 |
| D5M17 | Microbial<br>mat | 40-50 | Dome,<br>Site DA | Sep-2021 | Dry | 2851518 | 2706609 | 884 |
| D1M18 | Microbial<br>mat | 0-10 | Dome,<br>Site DA | Mar-2022 | Dry | 1666524 | 1593750 | 907 |
| D2M19 | Microbial<br>mat | 010-20 | Dome,<br>Site DA | Mar-2022 | Dry | 1308223 | 1264518 | 1022 |
| D3M20 | Microbial<br>mat | 20-30 | Dome,<br>Site DA | Mar-2022 | Dry | 1055947 | 1006567 | 893 |
| D4M21 | Microbial<br>mat | 30-40 | Dome,<br>Site DA | Mar-2022 | Dry | 687386 | 657723 | 800 |
| D5M22 | Microbial<br>mat | 40-50 | Dome,<br>Site DA | Mar-2022 | Dry | 1152282 | 1099173 | 863 |
| D1M23 | Microbial<br>mat | 0-10 | Dome,<br>Site DA | Mar-2023 | Dry | 9651131 | 8704738 | 1717 |
| C1M24 | Sediment | 0-10 | Circle<br>Site AD | Mar-2023 | Dry | 4077381 | 3607459 | 1277 |
| C1M25 | Sediment | 0-10 | Circle<br>Site AD | Mar-2023 | Dry | 4379044 | 3807033 | 1347 |
| S1 | Water | 0-10 | Churince | Nov-2012 | Not<br>reporte<br>d | 1659099 | 1384599 | 649 |
| S2 | Water | 0-10 | Churince | Nov-2012 | Not<br>reporte<br>d | 846704 | 593281 | 1070 |
| S3 | Water | 0-10 | Churince | Nov-2012 |  | 2033524 | 1476091 | 800 |
| S4 | Water | 0-10 | Churince | May-2013 | Not reported | 2170234 | 1771093 | 668 |
| S5 | Water | 0-10 | Churince | May-2013 | Not reported | 2448431 | 2155568 | 649 |
| S6 | Water | 0-10 | Churince | May-2013 | Not reported | 1634495 | 1388562 | 607 |
| S7 | Water | 0-10 | Churince | Oct-2013 | Not reported | 2379615 | 2072533 | 630 |
| S8 | Water | 0-10 | Churince | Oct-2013 | Not reported | 1762737 | 1509098 | 642 |
| S9 | Water | 0-10 | Churince | Oct-2013 | Not reported | 2715904 | 2328461 | 667 |
| S10 | Water | 0-10 | Churince | May-2014 | Not reported | 2028313 | 1711573 | 594 |
| S11 | Water | 0-10 | Churince | May-2014 | Not reported | 2192243 | 1941794 | 609 |
| S12 | Water | 0-10 | Churince | May-2014 | Not<br>reporte<br>d | 3012508 | 2544221 | 615 |

A heatmap of the core microbiome was generated using the plot_core function, which was implemented in microbiome package in R, specifying prevalence thresholds (0%, 10%, 20%, 30%, 40%, 50%, 60%, 70%, 80%, 90%, 100%) and detection thresholds (0.001, 0.003, 0.009). This approach provided a clear and detailed visualization of the core microbiome, facilitating the identification of present taxa and their relative prevalence in the analyzed samples.

The make_network function from the phyloseq package was used with a maximum distance value (max.dist) of 0.7 to define the connections between samples. The Bray-Curtis distance metric was used to calculate dissimilarity between samples, ranging from 0 (identical composition) to 1 (completely different composition), and considering the relative abundances of taxa.

### 2.7 Metagenome-assembled genomes (MAGs) of AD and Churince sites

We observed that the wetland at AD has been declining over the eight-year period that we have been studying it (due to poor management of this natural resource. Pérez Ortega 2020) leading to an apparent loss of microbiological diversity, including endemic microorganisms and of one of the most diverse archaeal communities in the world (Rodríguez-Cruz et al., 2024).

Here we compared the change in the number of metagenome-assembled genomes (MAGs) during this 8-year period of study. For this purpose, MAGs were assembled from the new metagenomic data obtained during 2023, using the binning tools MaxBin2 (v2.2.7) (Wu et al., 2015) with the options run_MaxBin.pl -thread 24 - contig file_scaffolds.fasta -out directory_maxbin -abund file_scaffolds.bam.abundance.txt, Metabat2 (v2:2.15) (Kang et al., 2019) with the options runMetaBat.sh -m 2500 file_scaffolds.fasta file_scaffolds.bam, and MetaCOAG (v1.0)(Mallawaarachchi & Lin, 2022) with the options MetaCoAG --assembler spades -- graph file_assembly_graph_with_scaffolds.gfa --contigs file_contigs.fasta --paths file_contigs.paths --abundance file_bam.abundance.txt --output directory_metacoag_out. We assessed the completeness and contamination of each MAG using the checkm (v1.1.3) (Parks et al., 2015)software.

The same procedure was applied to the Churince data, obtained from De Anda et al. (2018), which covers the period from 2012 to 2014 and includes 12 metagenomes. Given the number of MAGs analyzed in this study for the Archean Domes (912 MAGs), we only performed functional annotation for the Churince MAGs. For this, we used freely available hidden Markov model (HMM) databases for microbial metabolic genes of environmental/biogeochemical relevance. For example, we utilized the metabolic-hmms database (available at https://github.com/banfieldlab/metabolic-hmms), FOAM (Functional Ontology Assignments for Metagenomes) (Prestat et al., 2014), TIGRFAMS (Haft, 2003), and Pfam (Mistry et al., 2021)(V36.0). This annotation was then mapped to KEGG orthologs and normalized to the total number of coding sequences per genome.

A phylogenetic perspective of the MAGs obtained from the AD site (samples collected between 2016 and 2023) and El Churince (samples collected between 2012 and 2014) was inferred using the tree generated by GTDB-tk(Chaumeil et al., 2020) (v2.3.1) and visualized with iTOL (Letunic & Bork, 2021).

## 3.0 Results and discussion

### 3.1 Alpha and Beta Diversity Analysis for 16S rRNA Data from AD and PR

For the 16S tag data between the two Eastern sites: Pozas Rojas (PR) and Archean Domes (AD) with microbial mat (D) and orange circles (C), we had a total of 27 sequencing samples were processed, each starting with a variable number of initial reads. Filtering and denoising steps resulted in a substantial reduction in reads. After filtering, the number of reads per sample ranged from 29,152 in sample D6104 to 445,898 in sample D7110. Denoised reads also exhibited a gradual decrease compared to initial forward and reverse reads, reflecting the effectiveness of the applied cleaning steps.

Taxonomic analysis of the samples revealed significant diversity in sediment microbial phylum composition. Halobacterota emerged as the predominant phylum in several samples, reaching its highest abundance in sample C123 at 0.698. Bacteroidota was also highly represented, particularly in D6104 (0.495) and D8106 (0.333). Nanoarchaeota was notable in D6109 (0.264) and PR6 (0.158). Halanaerobiaeota was prominent in PR5 with an abundance of 0.212, while Firmicutes, although less dominant, had a notable presence in D10113 (0.152) and D8106 (0.140). Desulfobacterota also showed relevant abundances in D852 (0.174) and D953 (0.134).

Alpha diversity of the samples using 16S rRNA Sequence Data was assessed using the Observed richness (number of observed ASVs), Chao1 (species richness estimator), and Shannon Index (species diversity) estimates. The samples exhibited a wide range of species richness (see Table S1, S2, and S3 for more information on each sample, Summary statistics, encompassing read counts, and the number of unique ASVs are shown in Fig. 3A; see Fig. S1 for rarefication curves.

D7110 and D9112 samples showed the highest values of observed ASVs (1391 and 1330, respectively), indicating high species richness. In contrast, samples such as D7105 and D6104 had the lowest values (only 23 and 29, respectively), suggesting low species richness for those data points. This is confirmed by the Shannon Index, samples D7110 and D9112 again showed the highest Shannon index values (5.840 and 5.845, respectively), indicating high diversity of species. Conversely, samples D7105 and D6104 again had the lowest Shannon index values (2.612 and 2.575, respectively), suggesting low diversity and a less even distribution of species.

In AD, samples collected at 0-10 cm depth exhibited considerable variability in alpha diversity. Samples such as C123, C146 and C223, collected in 2023 and 2021 from the Orange Circles, showed relatively low Shannon values (2.808, 3.679, and 3.454, respectively). Nevertheless, some surface samples from PR, including PR1, PR2, and PR6, showed high Shannon values (5.376, 5.793, and 5.802, respectively), indicating that specific local factors may significantly influence surface microbial diversity (see Fig. S1) Intermediate depth samples --such as D7110 (10-20 cm) and D9112 (30-40 cm)--exhibited high species richness and diversity, with Shannon values of 5.840 and 5.845, respectively, indicative of diverse and balanced microbiological ecosystems. These soil layers may provide a more stable and less disturbed environment, favoring the formation and maintenance of diverse, balanced, and complex microbial communities. Deeper samples taken at 40-50 cm – i.e., C550 and D1155--also showed high diversity, with Shannon values of 4.519 and 4.966, respectively. These depths may include unique microhabitats and specific resources that promote high microbial diversity, possibly due to the accumulation of organic matter and less disturbance from external factors.

On the other hand, samples PR1, PR2, PR4, and PR6 exhibit a distinctive microbial profile that sets them apart from other groups in the NMDS analysis (Fig. 2). Their positive position on the MDS2 axis suggests that these microbial profiles are characterized by unique features not found in samples from orange circles (C) and Domes microbial mats (D)

**Figure 2.**
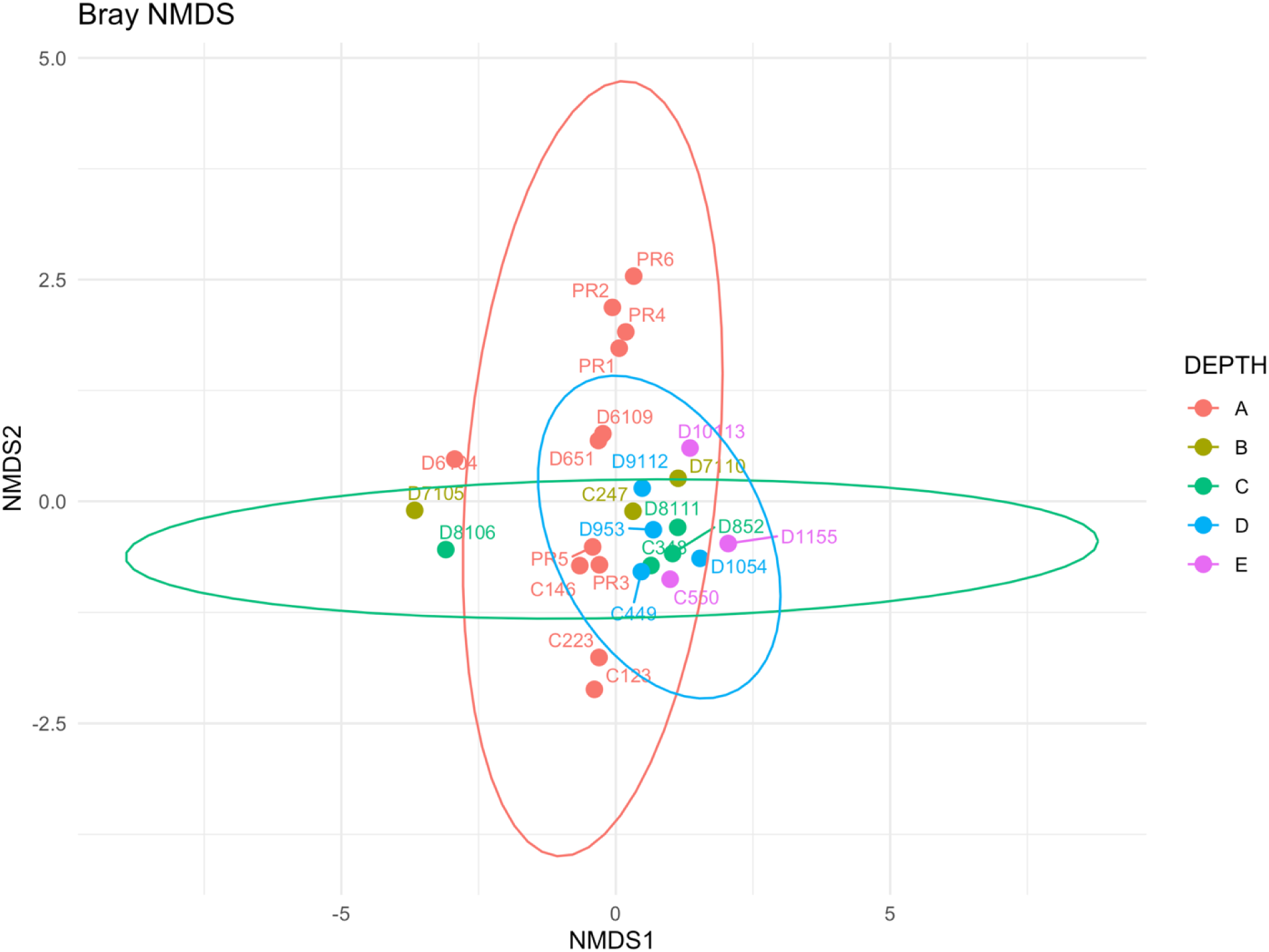
Non-metric multidimensional scaling (NMDS) ordination plot based on the Bray-Curtis dissimilarity matrix, illustrating the compositional dissimilarities between microbial communities across samples. The data were obtained from 16S rRNA gene sequencing of the samples. Each point represents a unique microbial community, with distances between points reflecting the degree of dissimilarity. Samples are color-coded according to their location to highlight spatial patterns in beta diversity. The color coding corresponds to sampling depths, where A = 0–10 cm, B = 10–20 cm, C = 20–30 cm, D = 30–40 cm, and E = 40–50 cm.

The total dispersion within the PR group indicates substantial variability in their microbiological composition. Samples PR3 and PR5 show relative proximity to some samples from group D (D953 and C449, taken at a depth of 30-40 cm) in the NMDS space (Fig.2), implying potential similarities in their microbial profiles. However, the differences between these groups remain pronounced enough to clearly distinguish between them. This may indicate that while groups PR and D share some microbial characteristics, they also exhibit significant differences in their composition, probably due to each site particular environmental filtering.

### 3.2 Core, shared and unique ASVs obtained using 16S rRNA Sequence Data in AD and PR

The distribution and overlap of ASVs between different sampling sites are depicted in Fig. 3 A. We found a high proportion of global (i.e., shared taxa in the three sites) core microbiota, that comprised 237 ASVs (see Fig. S2 for a detailed distribution of the shared ASVs between PR and AD). Of these, 215 ASVs were classified at the order level. Among these, the most predominant orders were Halobacterales, with 27 ASVs, representing approximately 11.39% of the classified ASVs; followed by Halanaerobiales, with 16 ASVs (6.75%), and Woesearchaeales, with 15 ASVs (6.33%). Additional orders with important representation include **Bacteroidales** with **12 ASVs** (**5.06%**), and **Phycisphaerales** and **Spirochaetales**, each with **10 ASVs** (**4.22%**). Together, these five orders account for **33.97%** of the classified ASVs. It is noteworthy that **22 ASVs** were not classified at the order level, which represents **9.28%** of the total ASVs. This finding highlights the need for improved taxonomic resolution to achieve a more comprehensive understanding of microbial diversity in the studied contexts. Orders such as **Actinomarinales**, **Alicyclobacillales**, **Anaerolineales**, and **Caulobacterales**, among others, are represented by a single ASV each, contributing only **0.42%** to the total classified ASVs. The shared ASVs among the three sites suggested the presence of a core microbial community that could be the product of connection (dispersion) among sites, involving microbes that are capable to adapt and survive in different physicochemical conditions. The ASVs data also highlight the notable number of unique ASVs in PR and D.

**Figure 3.**
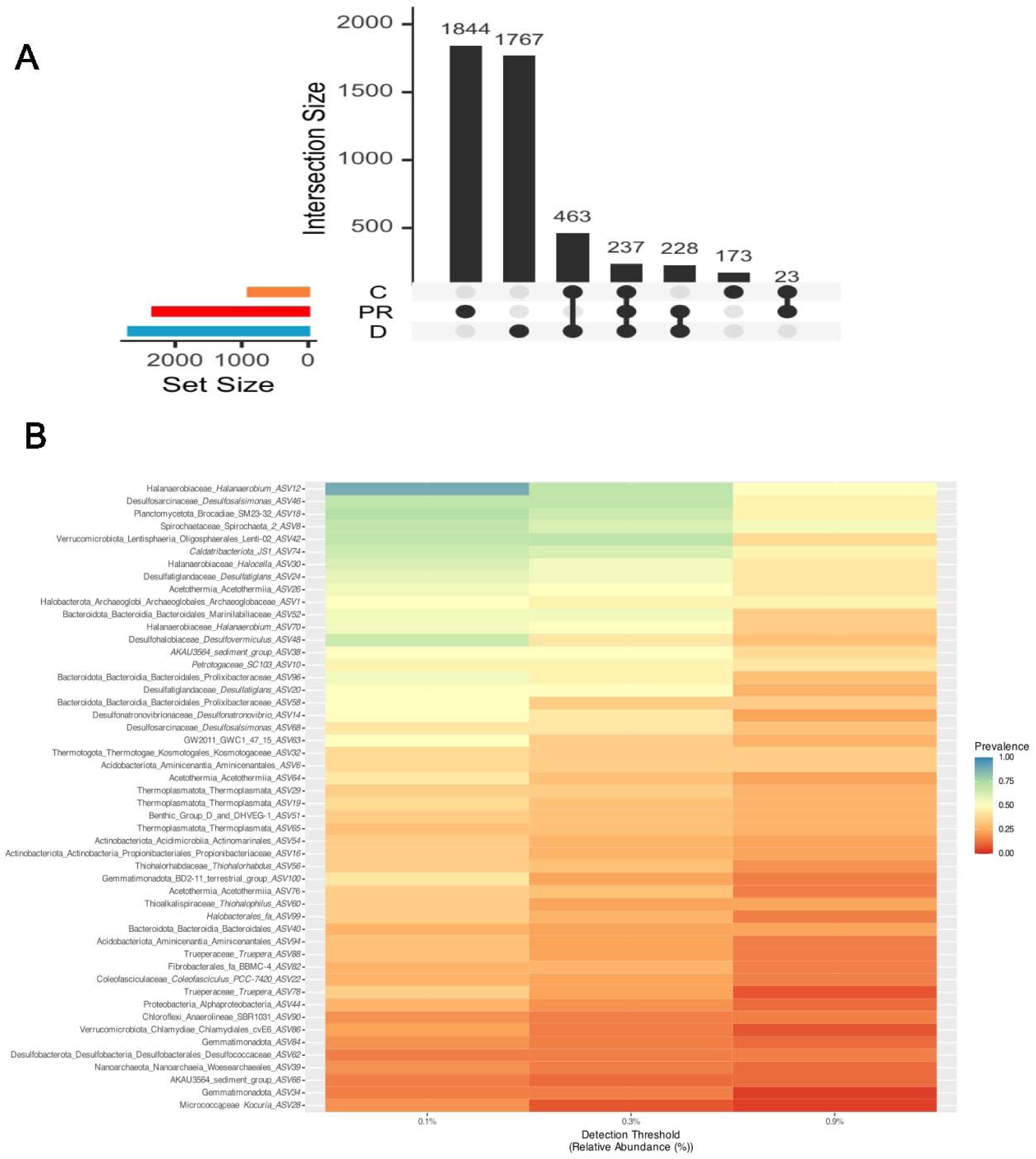
Core Microbiome Heatmap and Upset Plot Analysis from 16S rRNA gene sequencing samples. (A) Upset plot showing the shared and unique ASVs among different sampling sites where ‘C’ represents samples from the Circles at the Archean Domes, ‘D’ refers to other sampling points within the same site, and ‘PR’ corresponds to samples from the “Pozas Rojas” site. (B) Heatmap depicting the core microbiome from AD and PR sites, where detection of each taxon was calculated as the proportion of samples in which the relative abundance exceeded 1%. A prevalence threshold of 10% was applied to select the ASVs of interest.

The prevalence analysis of the ASVs in the samples, considering a minimum detection threshold of 0.1% relative abundance (Fig. 3 B), reveals pattern on the structure of the shared microbial community across the different study sites. The prevalence of ASVs was calculated as the proportion of samples in which each ASV was present above the detection threshold.

Accordingly, *Halanaerobium* ASV12 (see Fig. 3B) exhibited a notably high prevalence, being found in 88.46% of the samples, indicating its wide distribution and possible ecological relevance in these environments. Members of this genus have been found in high abundance in subterranean systems with high salinity, such as the Barnett Shale geological formation located in the Bend Arch-Fort Worth basin (USA), with relative abundances ranging from 17% to 33% (Liang et al., 2016a). Members of the genus *Halanaerobium* are known for carbohydrate fermentation and sulfur production through thiosulfate reduction (Ravot et al., 1997, 2005). Therefore, *Halanaerobium* is clearly an important genus that could play vital roles in organic matter biodegradation and sulfur production in fractured shale formations.

Another ASV with high prevalence among CCB ponds is *Desulfovermiculus* ASV48 (see Fig. 3B), that was detected in 65.38% samples. *Desulfovermiculus* spp. (Belyakova et al., 2006) has been reported as halophilic sulfate reducers, suggesting that sulfate reduction might contribute to sulfide production in high-salinity subterranean systems. The high prevalence of this genus among ponds suggests that this taxon is also an important component of the shared microbial community.

It is interesting to highlight that both *Halanaerobium* sp. and *Desulfovermiculus* sp. have been found in high-salinity environments and at shallow depths, such as in the Barnett Shale geological formation (Liang et al., 2016b), which could suggest that both *Halanaerobium* sp. and *Desulfovermiculus* sp. may be transported across different ponds from a deep aquifer. Nevertheless, sampling at greater depths, will be needed to analyze the microbial community within both the PR site and AD sites.

ASVs such as *Spirochaeta*_2 ASV8 and *Halocella* ASV30, with prevalences of 69.23%, also show a wide distribution among the samples, reinforcing the hypothesis of a set of common and predominant taxa in these sites. *Desulfosalsimonas* ASV46, has been described as a halophilic Gram-negative sulfate-reducing bacterium(Kjeldsen et al., 2010).Consequently, it is a common member of extreme hypersaline sediments, where salinities range from 120 to 270 NaCl l^-^ (Kjeldsen et al., 2010).In turn, *Desulfonatronovibrio* ASV14 has been described as an alkaliphilic, sulfate-reducing bacterium (ZHILINA et al., 1997)and it has been isolated from a soda-depositing lake (ZHILINA et al., 1997)*Desulfonatronovibrio* comprises motile vibrio bacteria that utilize only hydrogen and formate as electron donors, while using sulfate, sulfite, and thiosulfate, but not sulfur, as electron acceptors (ZHILINA et al., 1997). *Desulfonatronovibrio* ASV14 was found in 69.23% of the samples, suggesting to the possible involvement of these taxa in specific biogeochemical processes such as the sulfur biogeochemical cycle within the ponds analyzed in this study.

In contrast, some ASVs exhibited lower prevalence, such as *Kocuria* ASV28, which was detected in only 19.23% of the samples. This suggests that these taxa may occupy more restricted ecological niches or that their populations are more vulnerable to environmental fluctuations. Alternatively, it could indicate that the dispersal capacity of this bacterium is limited, hindering its ability to spread across the different ponds within the Cuatro Ciénegas Basin.

### 3.3 Core, shared and unique microbial taxa between AD sites and Churince using whole genome metagenome data

Overall, alpha diversity values measured using the Shannon index for samples from the Archaean Domes site, collected between 2016 and 2023 at depths ranging from 0 to 50 cm, exhibited considerable variability. The dataset covers a seven-year period with samples collected at various depths: 0-10 cm (depth A), 10-20 cm (depth B), 20-30 cm (depth C), 30-40 cm (depth D), and 40-50 cm (depth E).

For samples collected at depth A (0-10 cm) at the site Archaean Domes, Shannon index values ranged from a minimum of 4.40 in 2021 (D1M13) to a maximum of 6.23 in 2018 (D1M04). Notably high values were observed in 2020 (D1M07) and 2022 (D1M18), both with a Shannon index of 5.92. Lower diversity values were recorded in 2017 (D1M03) with an index of 5.12 and in 2021 (D1M13) with an index of 4.40.

At depth B (10-20 cm), Shannon index values were 6.77 for the sample collected in 2021 (D2M14) and 6.04 for the sample collected in 2022 (D2M19). At depth C (20-30 cm), Shannon index values for samples such as D3M15 (2021) and D3M20 (2022) were 6.26 and 6.03, respectively. For samples from depth D (30-40 cm), the Shannon index values included 6.54 for 2020 (D4M11) and 6.55 for 2021 (D4M16). At depth E (40-50 cm), values ranged from 6.23 in 2020 (D5M12) to 6.60 in 2021 (D5M17) and 6.47 in 2022 (D5M22).

In contrast, samples from the CH site, collected between 2012 and 2014, demonstrated consistently higher Shannon index values. For depth A (0-10 cm), Shannon index values ranged from 5.75 to 7.04, with notable indices of 7.04 for sample S1 in 2012 and 6.76 for sample S10 in 2014

The prevalence analysis of taxa in the AD and Churince samples, using a minimum detection threshold of 0.1% relative abundance, revealed a set of taxa consistently detected across a large fraction of the samples (Fig. 4A). This suggests that these organisms are capable of dispersing among the different ponds in the CCB and may play fundamental roles in the studied microbial ecosystems. Specifically, the shared microbiome between the studied sites includes approximately 21.2% Archaea, represented by taxa such as *Halodesulfurarchaeum formicicum*, *Halorubrum*, *Halorhodospira halophila*, *Haloterrigena* spp., *Halovivax asiaticus*, *Salinarchaeum* sp. Harcht-Bsk1, and *Thermoplasmata* archaeon. In contrast, 78.8% of the taxa in this shared microbiome are Bacteria, encompassing a diverse range of groups including Desulfobacteraceae (e.g., *Desulfosalsimonas propionicica*, *Desulfatitalea tepidiphila*, *Desulfonatronovibrio magnus*, *Desulfonema ishimotonii*, *Desulfohalobium retbaense*), Deltaproteobacteria (e.g., *Deltaproteobacteria bacterium*), and Spirochaetaceae (e.g., *Spirochaetaceae bacterium*), as well as other taxa such as Actinobacteria, Acidobacteria, Gemmatimonadetes, Halanaerobium, Halomicrobium, Halomonas, Halorussus, Mariniphaga, Nitrospirae, Phycisphaeraceae, Planctomycetes, Puniceicoccaceae, Rhodobacteraceae, Rhodosalinus, and *Candidatus Melainabacteria*.

**Figure 4.**
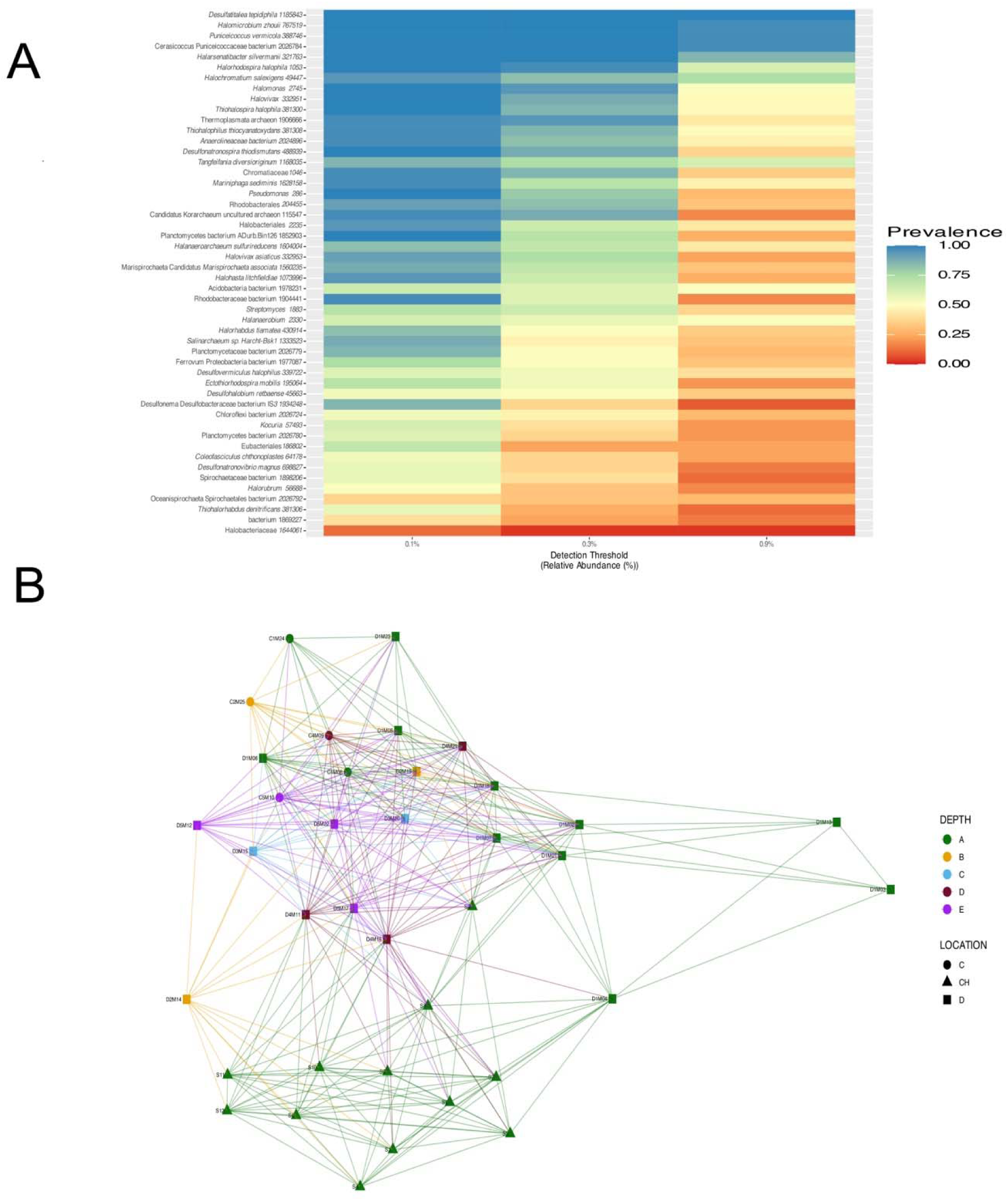
Heatmap of the microbiome from the AD site and Churince, alongside a network analysis based on whole metagenome sequencing data. In the figure, the sites are labeled as follows: ‘D’ for Domes and ‘C’ for Circles, both located within the AD site, and ‘CH’ for Churince (samples S1 to S12). (A) Heatmap depicting the core microbiome. Prevalence thresholds were set at 0%, 10%, 20%, 30%, 40%, 50%, 60%, 70%, 80%, 90%, and 100%, while detection thresholds were set at 0.001, 0.003, and 0.009. (B) Network analysis of the same metagenomic dataset transformed to relative abundance, illustrating the relationships between samples based on taxon composition. The make_network function from the phyloseq package was used with a maximum distance value (max.dist) of 0.7 to define connections between samples. Bray-Curtis distance metric was applied to calculate dissimilarity between samples, ranging from 0 (identical composition) to 1 (completely different composition), considering the relative abundances of taxa. The color coding corresponds to sampling depths, where A = 0–10 cm, B = 10–20 cm, C = 20–30 cm, D = 30–40 cm, and E = 40–50 cm.

Taxa with high prevalence, such as *Halanaerobium* sp. (63%), *Desulfovermiculus halophilus* (60%) and Spirochaetaceae bacterium 1898206, 57%), although they are not present in all samples, its high prevalence suggests that they can disperse among the different aquatic systems, and that they may play an important role in the microbial communities, indicating environmental stability and communication between communities that share the same deep aquifer. This is particularly important for the rare biosphere, that is why we lowered the detection values used in the analysis were 0.001, 0.003, and 0.009, representing permissive thresholds of relative abundance. These thresholds help identify taxa that are present even in low proportions, providing a more inclusive view of microbial diversity.

To explore the relationships between samples based on taxon composition A network analysis was performed using the metagenomic dataset transformed to relative abundance. For instance, a connection between sample D4M11 (AD depth 30-40 cm), and sample S8 (Churince), was found, as well as the connections between metagenomes D5M17 (AD depth 40-50 cm) and S8 (Churince) (Fig. 4), again suggesting a possible connection between the deep aquifer and these two geographic sites, in CCB despite their geographic distance as well as nearly 10 years between samplings. This is very interesting since the deep aquifer seems to work as a vault that preserved Churince diversity despite the demise of the system. This is even more evident the deeper the sample. At the depth of 30-40 cm at AD could be under stronger influence of the deep aquifer, allowing the dispersal of microorganisms that are also found in Churince.

Previous isotopic studies have shown that groundwater from the deep aquifer is the main source water for all the different CCB aquatic systems (Wolaver et al., 2013). We suggest that this deep aquifer has preserved bacterial lineages by conserving similar conditions to the ancient ocean maintaining these ancient microbial lineages isolated from their marine relatives for millions of years, time which has been sufficient to allow the emergence of a high microbial diversity. Also, we consider that this is the source of the shared microbial species that we find in the different aquatic systems in CCB.

Nevertheless, we agree that we will need futures analyses involving the generation of new metagenomic data of deeper samples along the different site in CCB Sadly, in some systems this will not be possible, as for instance all the area of Churince and nearby sites in the west wing of CCB are all dry now. Due to this unfortunate loss, we decided to explicitly compare the number of MAGs that have been lost over just 8 years of studying the AD site with the concern that the site might dry up similarly to Churince.

### 3.4 MAGs from 2016 to 2023 show a decrease of microbial diversity in Archean Domes site

From the 25 metagenomes generated between 2016 and 2023 for AD site, a total of 912 MAGs (171 from the domain Archaea and 647 from Bacteria) with a completeness value > 20% were assembled (see Fig. 5, panels A and B for a phylogenetic perspective). Our study identified 70 taxonomic classes for bacterial MAGs (see Fig. S3 and table 3) with a completeness value > 20% and then were used to reconstruct a phylogeny (Fig. 5 A and B). On the other side, for the Archaea, our study identified 13 taxonomic classes (see Fig. S3).

**Table 3.**
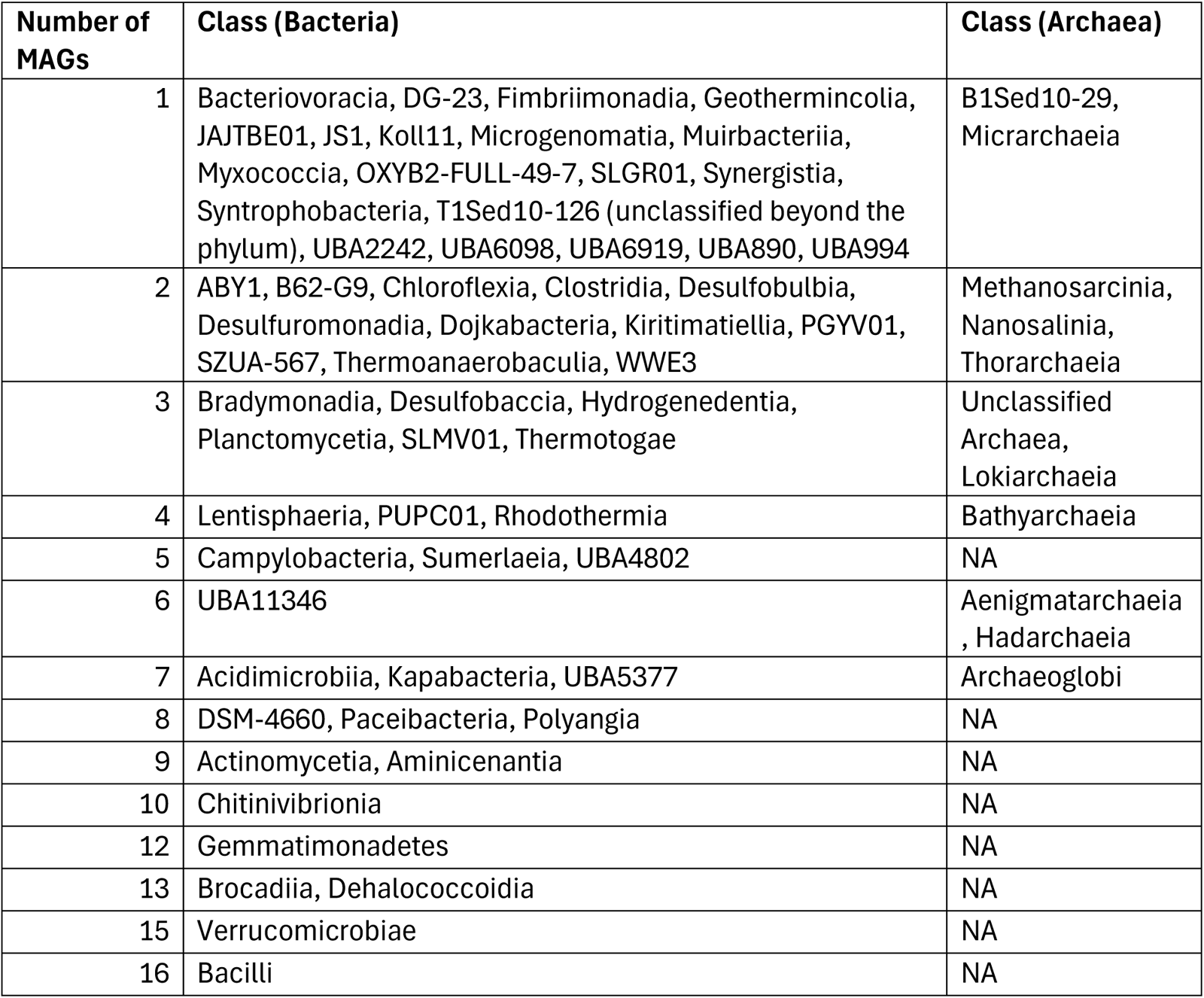
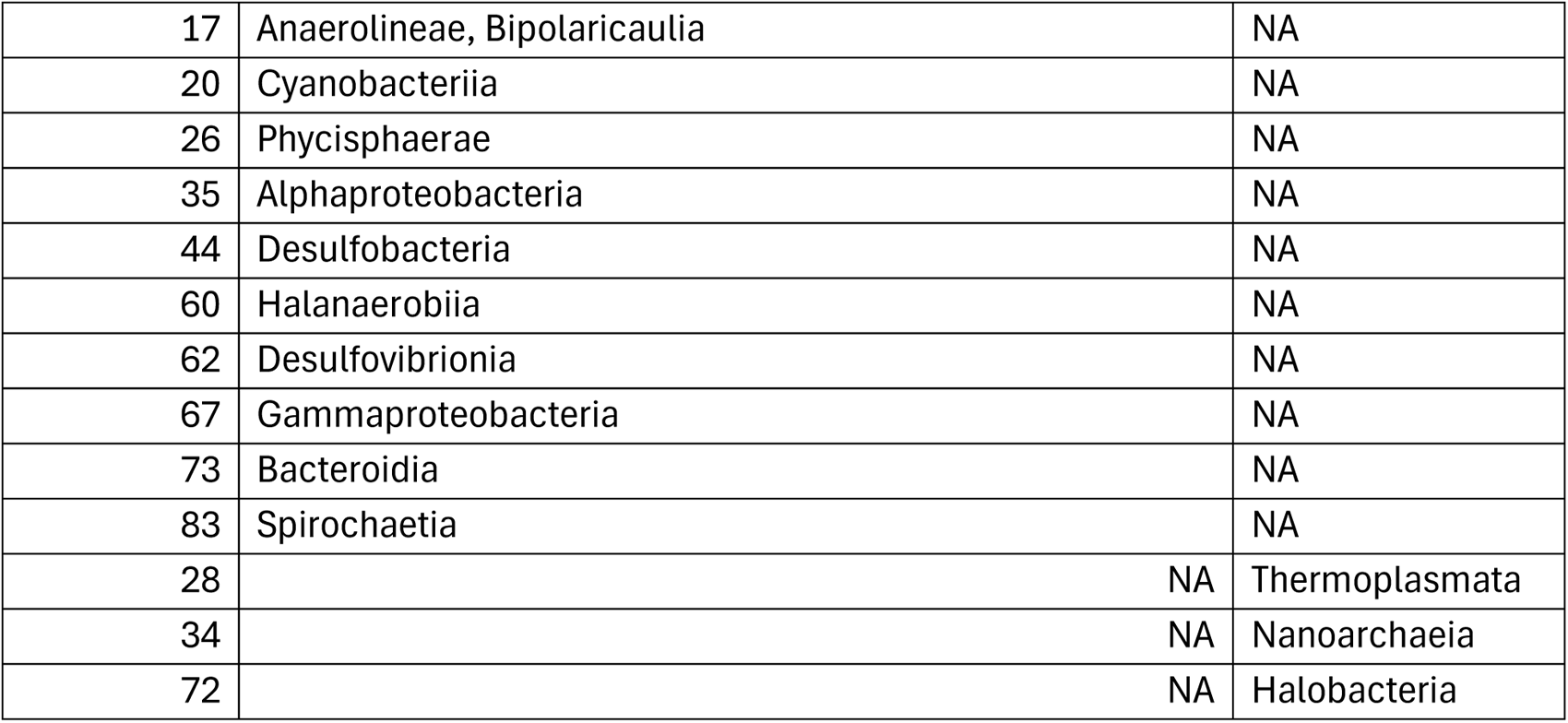
MAGs from AD site from years 2016 to 2023.

**Figure 5.**
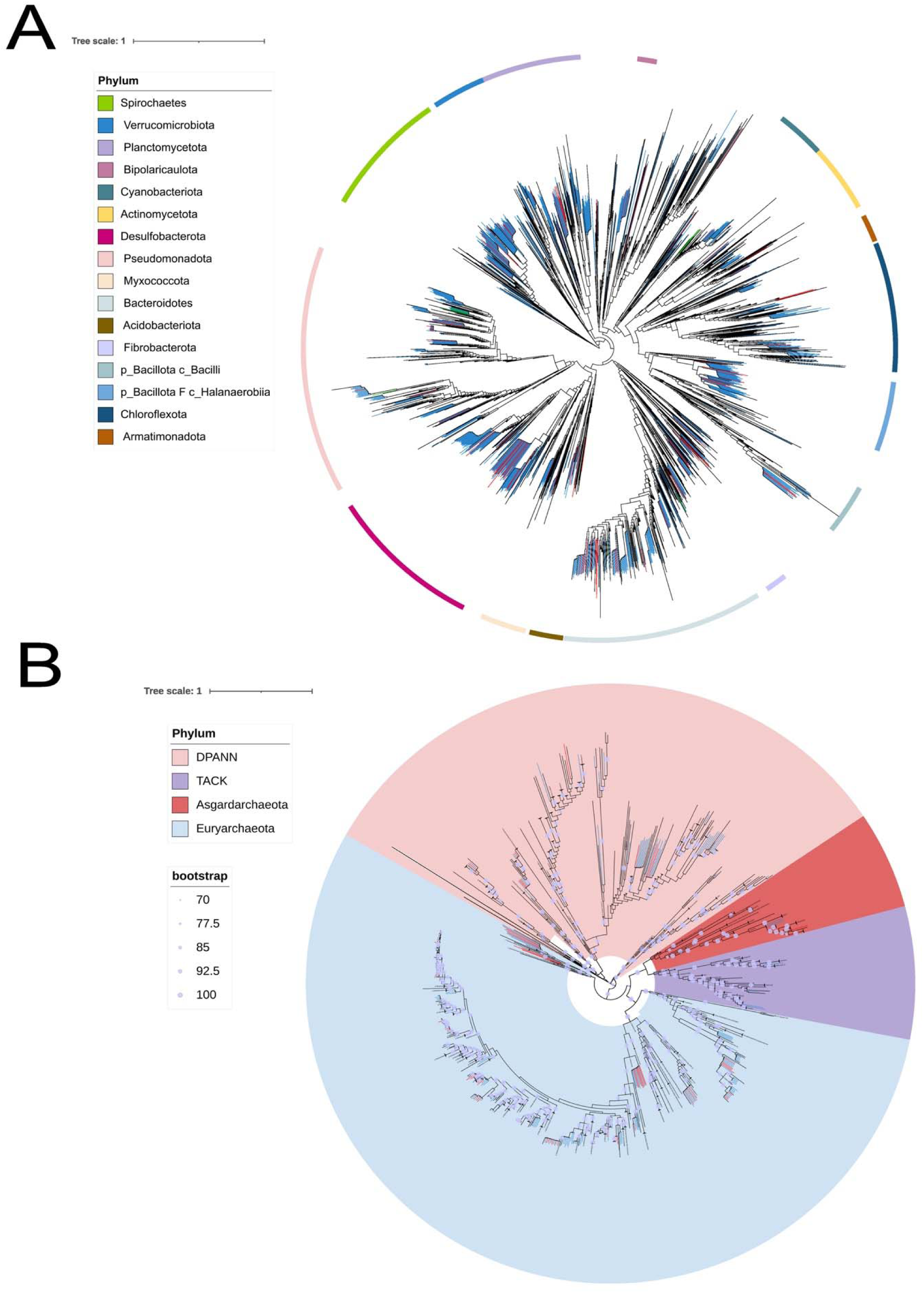
Phylogenetic analysis of MAGs from the AD and Churince sites (2016–2023). The phylogenetic tree was constructed using GTDB-tk v2.3.1 and visualized in iTOL. Red branches indicate MAGs from the AD site assembled in 2023, blue branches correspond to MAGs from the same site assembled between 2016 and 2022, and green branches represent MAGs from the Churince site

These results underscore the taxonomic diversity of archaea in our dataset, with a notable predominance of MAGs belonging to the Halobacteria class.

Sadly, we observed a declining trend in the number of MAGs recovered as the samples become more recent (see Fig. S3B). Specifically, the oldest sample, D1M01, had 125 MAGs, while the most recent sample, D1M23, contains only 22 MAGs. This decrease is evident in several intermediate samples (see Fig. S3B): D1M01 (125 MAGs), D1M02 (34 MAGs), D1M03 (46 MAGs), D1M04 (47 MAGs), D1M05 (91 MAGs), D1M06 (92 MAGs), D1M07 (89 MAGs), D1M13 (39 MAGs), D1M18 (8 MAGs), and D1M23 (22 MAGs). Examining these data series, we note that after sample D1M01, which has the highest number of MAGs, there is a significant drop in sample D1M02, which has only 34 MAGs. This trend remains relatively constant with minor fluctuations until reaching sample D1M18, which has the lowest number of MAGs at just 8. Although there is a slight increase in sample D1M23 (22 MAGs) compared to D1M18, it is still significantly lower than in the older samples.

These data series suggest a potential loss of microbiological diversity in our samples over time (Fig. 6). The observed fluctuations in the intermediate samples (D1M05, D1M06, D1M07) may indicate specific events that temporarily affected microbiological diversity, but the overall trend points to a decline. The loss of diversity observed in the more recent samples from AD may reflect changes in the connectivity of the deep aquifer with the surface layers, possibly due to environmental or anthropogenic factors. This is consistent with the drying of the Churince system, the reduction of the water levels in many systems of CCB and the latent risk of a similar event occurring at all the other CCB sites, including PR, our actual only safe site.

**Figure 6.**
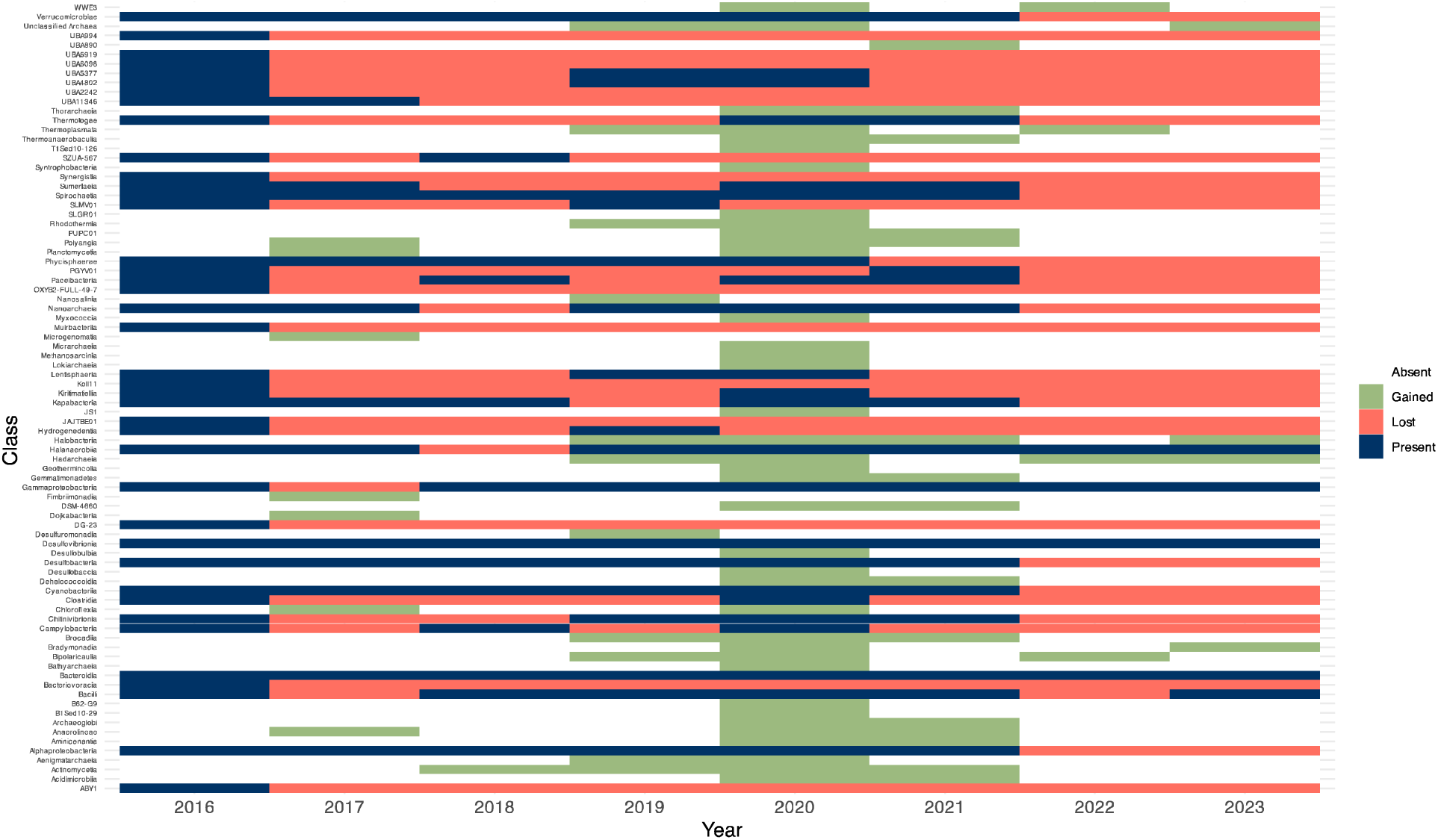
This heatmap compares the taxonomic classes gained or lost over the sampling period from 2016 to 2023 in the AD site. A taxonomic class is considered present if at least one MAG of that class is detected from the first year of sampling (2016). If a MAG from a present class is not found in the following year, it is considered a loss. Conversely, if a MAG from a class is not present in 2016 but appears in any subsequent year, it is considered a gain. Rows represent taxonomic classes, and columns represent the sampling years. Cells are colored according to the presence, loss, or gain of MAGs in different taxonomic classes over time.

The Pearson correlation coefficient was -0.1126 (p > 0.05), suggesting that there is no significant linear relationship between the proportion of MAGs by taxonomical class and the average sequencing depth. This result indicates that variations in the proportion of MAGs by class are not linearly associated with sequencing depths. In contrast, the Spearman correlation coefficient was -0.4517 (p < 0.05), revealing a moderate negative monotonic relationship. This suggests that, in general, the average sequencing depths tend to be lower when the proportion of certain microbial classifications is high, and vice versa.

As mentioned in the Methods section, we attempted to assemble MAGs from the 12 metagenomes obtained between 2012 and 2014 from the El Churince site. Unfortunately, we were only able to recover 5 MAGs with more than 20% completeness and less than 10% contamination, all of which belong to the Bacteria domain. These MAGs are listed in Table 4.

**Table 4.**
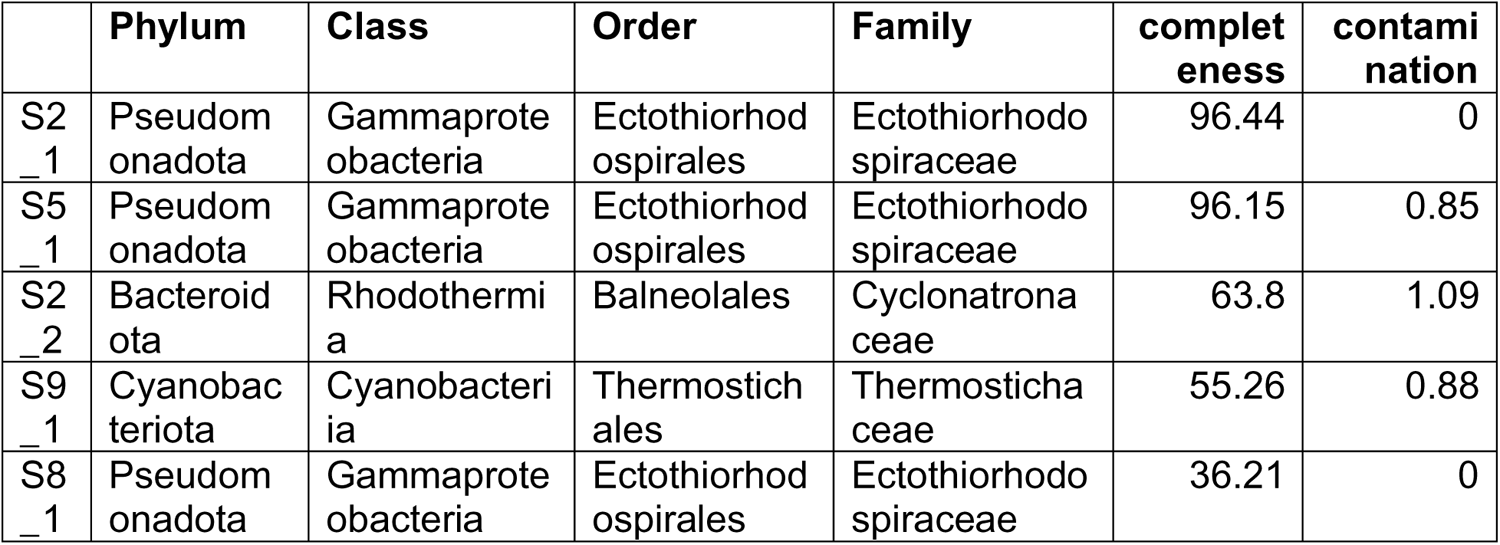
MAGs from the Churince site. This table lists MAGs with completeness values greater than 20% and contamination values under 10%.

As shown in Table 4, we recovered the largest number of MAGs (3 out of 5) corresponding to the genus *Ectothiorhodospira* at the Churince site. Interestingly, at the Archean Domes site, we also assembled 4 MAGs from the genus *Ectothiorhodospira*. When comparing the average nucleotide identity (ANI) between two MAGs of *Ectothiorhodospira* from each site (Archean Domes and Churince; see Fig. S4A), focusing on those with the highest completeness and lowest contamination values (93.9% completeness and 1.42% contamination for MAG M08_12 from the AD site), we observed an ANI of 90.49%. According to the commonly accepted threshold of 95-96% similarity to consider two genomes as belonging to the same species (Richter & Rosselló-Móra, 2009), this suggests that MAGs C1M08_12 and S2_1 may represent different species within the *Ectothiorhodospira* genus. However, since we are not comparing complete genomes (C1M08_12 has 93.9% completeness vs. 96.44% completeness for S2_1), we cannot entirely rule out the possibility that these two MAGs belong to the same species. Furthermore, when exploring the metabolic capabilities of both MAGs (see Fig. S4B), we found no notable differences between their metabolisms (see Fig. S4B for details). In terms of the metabolic capabilities of MAG S2_1 from Churince, functional annotation revealed genes associated with assimilatory sulfate reduction, including PAPSS (3’-phosphoadenosine 5’-phosphosulfate synthase) (Peng & Verma 1995), sat (sulfate adenylyltransferase) (Carlson et al. 2021), cysNC (bifunctional enzyme CysN/CysC), cysH (phosphoadenosine phosphosulfate reductase), and cysJ (sulfite reductase [NADPH] flavoprotein alpha-component) (Hummerjohann et al. 1998). Additionally, we identified the soxB, soxC, and soxY genes as part of the sulfur oxidation (Sox) multienzyme complex (Ghosh, Mallick, & DasGupta 2009), which is pivotal in the oxidation of reduced sulfur compounds like thiosulfate to sulfate. The presence of this complex indicates that this organism probably takes place in sulfur oxidation, a key process in energy extraction and sulfur cycling within sulfur-rich environments such as been reported in CCB (De Anda et al., 2018).

We also detected genes involved in dissimilatory sulfate reduction, such as ***aprAB*** (adenylylsulfate reductase) and ***dsrA*** (dissimilatory sulfite reductase alpha subunit) (Simon & Kroneck 2013). In addition to sulfur metabolism, several genes linked to nitrogen metabolism were found. These include nitrogen fixation genes ***nifD*** and **nifK**, encoding nitrogenase molybdenum-iron proteins critical for the conversion of atmospheric nitrogen into ammonia (Fani, Gallo & Liò 2000). Genes like ***nirS*** and ***norB*** (Black et al. 2016) were also present, playing essential roles in denitrification by reducing nitrite to nitric oxide and subsequently to nitrogen gas. Furthermore, the detection of ***nosZ*** (Black et al. 2016), responsible for reducing nitrous oxide to nitrogen, and ***amoA*** (Li et al. 2015), involved in ammonia oxidation during nitrification, suggests a comprehensive capacity for nitrogen cycling within this MAG.

## 4.0 Concluding remarks

During more than 20 years of molecular ecology been used to describe CCB diversity, we have felt all the time the need to describe as much as possible before it disappeared. Following that logic, we devoted 16 years to study Churince. As a result, this is probably the best know ecosystem in Mexico (see Springer series of 6 books; Souza & Eguiarte, 2018-2022). In this last work, putting all together all our results, we can highlight the presence of common microorganisms in the different sites. For instance, *Desulfovibrio* spp. is found both in the microbial community shared between the PR and AD, and between Churince and AD sites. Additionally, in the comparison of MAGs that persisted over the eight years of sampling, *Desulfovibrio* is the only taxonomic class of MAGs that remained prevalent throughout this period. These sulfate-reducing microorganisms have been reported from granitic groundwater sampled at a depth of 50-600 meters, frequently occur in deep granitic rock aquifers (Pedersen, 1997; 2000), which may support our hypothesis of microorganism transport from the deep aquifer to various surface ponds such as Pozas Rojas, AD, and Churince. The substantial presence of *Ectothiorhodospira* MAGs at multiple sites such as AD and Churince and their potential differences in species classification underscore the diversity within this genus. Furthermore, the persistence of *Ectothiorhodospira* sp. at both sites could suggest that there is also a transport of microbiological diversity from the deep aquifer mantle to different ponds within CCB, or it could indicate that the physicochemical conditions are maintaining suitable conditions for the proliferation of these microorganisms.

All these results give us a slight amount of hope, that if we protect Pozas Rojas there is a vault that is small but diverse and can preserve the dynamics between the deep aquifer and surface water. To this day, the ranchers around PR have donated water to the ecosystem thanks to the constant work of environmental education by us at first and then by “Plan 2040 (https://plan-2040.org/), the actual owners of the site”.

Another hopeful news is that more than 50 deep wells have been cancelled by the federal government in 2024 due to violation to water permits. We still working to change the way agriculture works in the desert, but that is a long-haul fight.

## 6.0 Acknowledgments

Ulises Erick Rodriguez Cruz is a doctoral student from the Programa de Doctorado en Ciencias Biomédicas, Universidad Nacional Autónoma de México (UNAM) and has received CONAHCYT fellowship 857544. Manuel Ochoa-Sánchez is a doctoral student from Posgrado en Ciencias Biológicas, Universidad Nacional Autonómas de México (UNAM) and acknowledges a CONAHCYT fellowship (CVU: 917392). We would like to thank Dra. Rosalinda Tapia-Lopez, Dra. Erika Aguirre-Planter and Manuel Rosas from the Instituto de Ecología, Universidad Nacional Autónoma de México, for technical and field assistance. We also thank PRONATURA Noreste for the access to the Pozas Azules ranch, and to Dra. Eria Rebollar-Caudillo and Dr. Arturo Becerra-Bracho for their valuable feedback on the manuscript. We also want to thank Rodrigo Garcia-Herrera, for facilitating the use of the high-performance computing cluster “Patung” located on Laboratorio Nacional de Ciencias de la Sostenibilidad (LANCIS), Instituto de Ecología (UNAM).

## 8.0 Supplementary tables

**Table S1.** Alpha diversity values for each sample.

|  | observed | chao1 | diversity_shannon |
| --- | --- | --- | --- |
| C123 | 31 | 31 | 2.808301 |
| C146 | 140 | 140 | 3.678725 |
| C223 | 54 | 54 | 3.454122 |
| C247 | 365 | 365 | 4.782714 |
| C348 | 459 | 459 | 4.975523 |
| C449 | 340 | 340 | 4.718458 |
| C550 | 326 | 326 | 4.518888 |
| D10113 | 238 | 238 | 4.018886 |
| D1054 | 574 | 574 | 5.244115 |
| D1155 | 492 | 492 | 4.965619 |
| D6104 | 29 | 29 | 2.574896 |
| D6109 | 462 | 462 | 4.817216 |
| D651 | 395 | 395 | 4.598852 |
| D7105 | 23 | 23 | 2.61211 |
| D7110 | 1391 | 1391 | 5.840409 |
| D8106 | 51 | 51 | 3.046637 |
| D8111 | 780 | 780 | 4.636291 |
| D852 | 337 | 337 | 4.599051 |
| D9112 | 1330 | 1330 | 5.845641 |
| D953 | 585 | 585 | 5.291147 |
| PR1 | 757 | 757 | 5.375644 |
| PR2 | 1078 | 1078 | 5.793846 |
| PR3 | 159 | 159 | 4.019634 |
| PR4 | 910 | 910 | 5.620532 |
| PR5 | 185 | 185 | 3.898878 |
| PR6 | 1083 | 1083 | 5.802379 |

**Table S2.** Reads summary from 16S rRNA data.

|  | input | filtered | denoised<br>F | denoised<br>R | merged | nonchim |
| --- | --- | --- | --- | --- | --- | --- |
| C123 | 5142 | 4689 | 4373 | 4298 | 3241 | 2882 |
| C146 | 147232 | 130409 | 126785 | 126651 | 106156 | 81928 |
| C223 | 5370 | 5010 | 4610 | 4526 | 3839 | 3824 |
| C247 | 120352 | 105763 | 101266 | 101011 | 84309 | 77474 |
| C348 | 176956 | 156267 | 150596 | 150269 | 128679 | 119121 |
| C449 | 149585 | 131642 | 126695 | 125889 | 106917 | 96401 |
| C550 | 124611 | 112666 | 107817 | 106700 | 92450 | 83795 |
| D10113 | 1036778 | 83431 | 78641 | 77912 | 68131 | 54553 |
| D1054 | 170113 | 152637 | 146144 | 145269 | 127516 | 119877 |
| D1155 | 205964 | 183580 | 176817 | 176217 | 151911 | 134529 |
| D6104 | 312427 | 29152 | 24967 | 24289 | 18432 | 10437 |
| D6109 | 255831 | 189418 | 178198 | 176546 | 153817 | 133267 |
| D651 | 182483 | 164905 | 159361 | 159006 | 138795 | 123673 |
| D7105 | 255096 | 34071 | 30773 | 29744 | 23997 | 10853 |
| D7110 | 478005 | 445898 | 436332 | 436135 | 410732 | 400983 |
| D8106 | 612648 | 154351 | 146421 | 143638 | 124845 | 60431 |
| D8111 | 481564 | 422719 | 405143 | 403220 | 346952 | 320726 |
| D852 | 99059 | 86444 | 81849 | 81405 | 70186 | 66055 |
| D9112 | 455572 | 420495 | 407117 | 405873 | 376885 | 368009 |
| D953 | 188739 | 165668 | 158756 | 157386 | 133676 | 124271 |
| PR1 | 243765 | 218115 | 207992 | 207847 | 175115 | 156389 |
| PR2 | 248008 | 220087 | 208585 | 207821 | 183829 | 176696 |
| PR3 | 100676 | 91188 | 88456 | 88481 | 76391 | 64736 |
| PR4 | 226901 | 205019 | 194245 | 193919 | 166780 | 155200 |
| PR5 | 178202 | 160080 | 155695 | 155784 | 130577 | 99353 |
| PR6 | 214900 | 193180 | 181734 | 180747 | 154470 | 144811 |

**Table S3.** Informaton for samples used in the analisys of 16S rRNA metabarcoding data.

|  | LOCATION | PRIMER | cm in deep | SITE | sampling year |
| --- | --- | --- | --- | --- | --- |
| C146 | C | 515F | 0-10 | ORANGE-CIRCLE | 2020 |
| C247 | C | 515F | 10-20 | ORANGE-CIRCLE | 2020 |
| C348 | C | 515F | 20-30 | ORANGE-CIRCLE | 2020 |
| C449 | C | 515F | 30-40 | ORANGE-CIRCLE | 2020 |
| C550 | C | 515F | 40-50 | ORANGE-CIRCLE | 2020 |
| D651 | D | 515F | 0-10 | DOME | 2020 |
| D852 | D | 515F | 10-20 | DOME | 2020 |
| D953 | D | 515F | 20-30 | DOME | 2020 |
| D1054 | D | 515F | 30-40 | DOME | 2020 |
| D1155 | D | 515F | 40-50 | DOME | 2020 |
| D6109 | D | 515F | 0-10 | DOME | 2021 |
| D7110 | D | 515F | 10-20 | DOME | 2021 |
| D8111 | D | 515F | 20-30 | DOME | 2021 |
| D9112 | D | 515F | 30-40 | DOME | 2021 |
| D10113 | D | 515F | 40-50 | DOME | 2021 |
| PR1 | PR | 515F | 0-10 | RED-POND | 2020 |
| PR2 | PR | 515F | 0-10 | RED-POND | 2020 |
| PR3 | PR | 515F | 0-10 | RED-POND | 2020 |
| PR4 | PR | 515F | 0-10 | RED-POND | 2020 |
| PR5 | PR | 515F | 0-10 | RED-POND | 2020 |
| PR6 | PR | 515F | 0-10 | RED-POND | 2020 |
| PR7 | PR | 515F | 0-10 | RED-POND | 2020 |
| PR9 | PR | 515F | 0-10 | RED-POND | 2020 |
| C123 | C | 515F | 0-10 | ORANGE-CIRCLE | 2023 |
| C223 | C | 515F | 10-20 | ORANGE-CIRCLE | 2023 |
| D7105 | D | 515F | 0-10 | DOME | 2021 |
| D6104 | D | 515F | 0-10 | DOME | 2021 |

---

**Figure S1.**
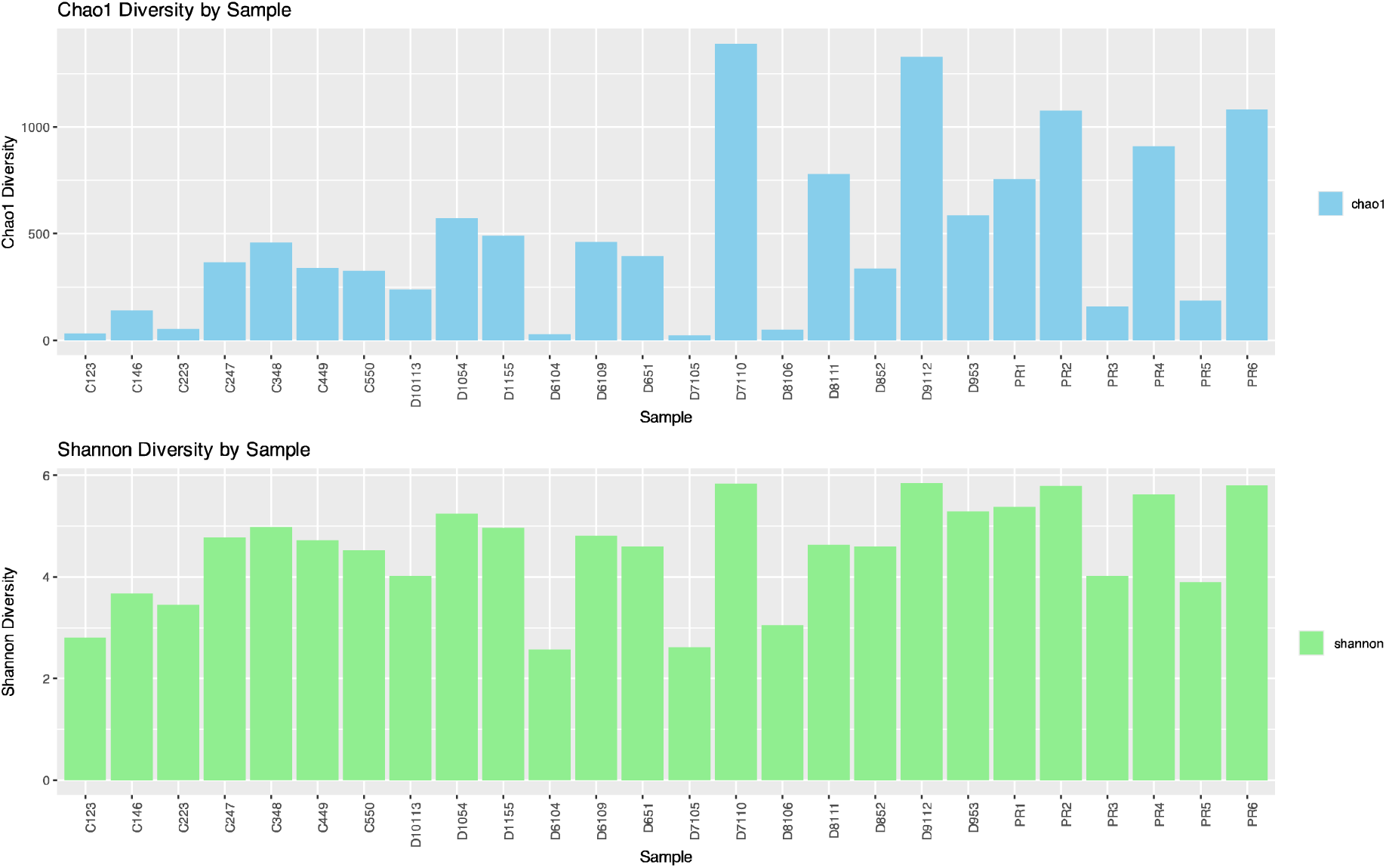
Alpha diversity indices for each sample, measured as (A) Chao1 and (B) Shannon indexes

**Figure S2.**
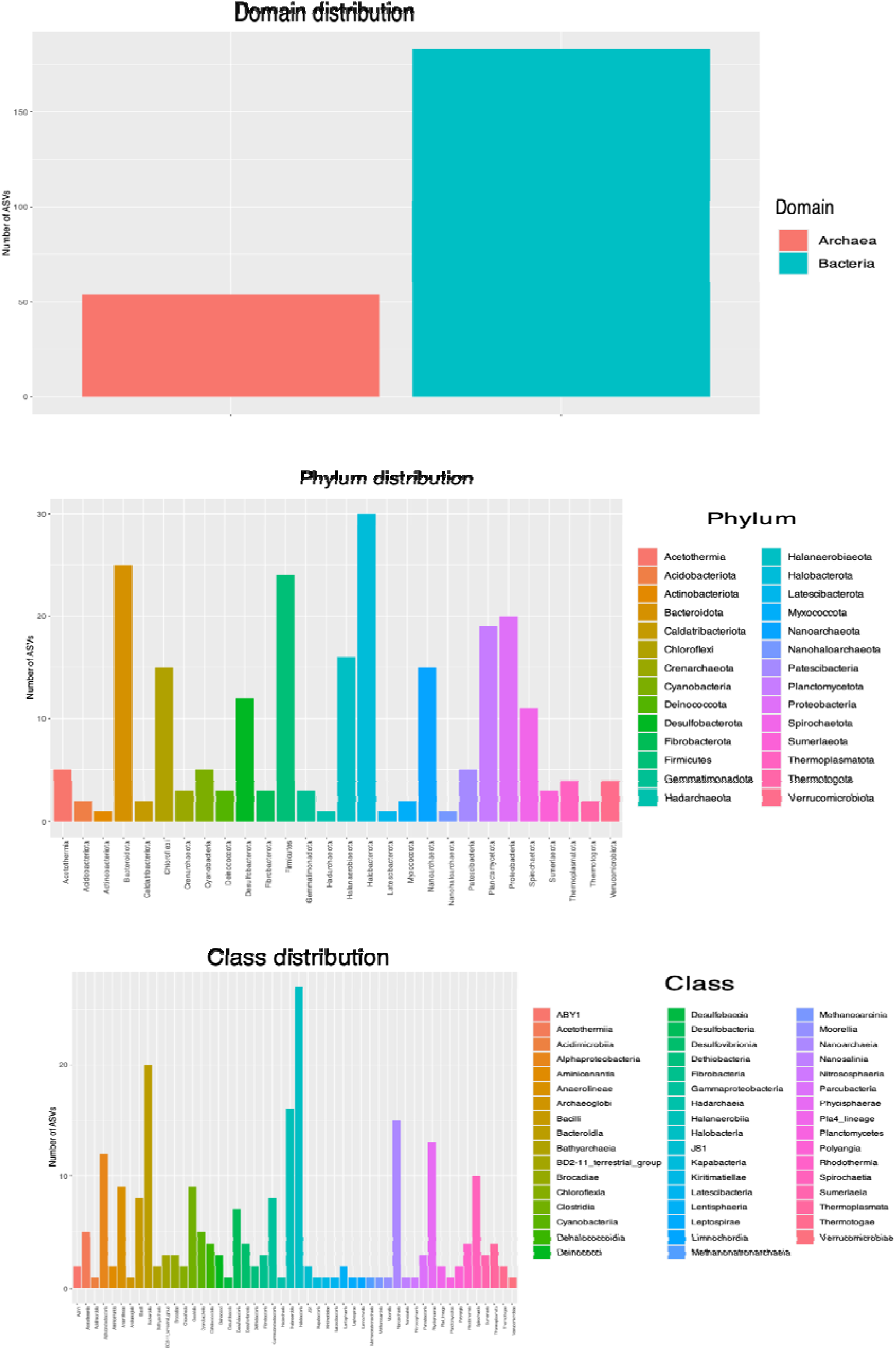
Distribution of shared ASVs between the AD and PR sites. (A) Classification of ASVs at the domain level. (B) Classification of ASVs at the phylum level. (C) Classification of ASVs at the class level.

**Figure S3.**
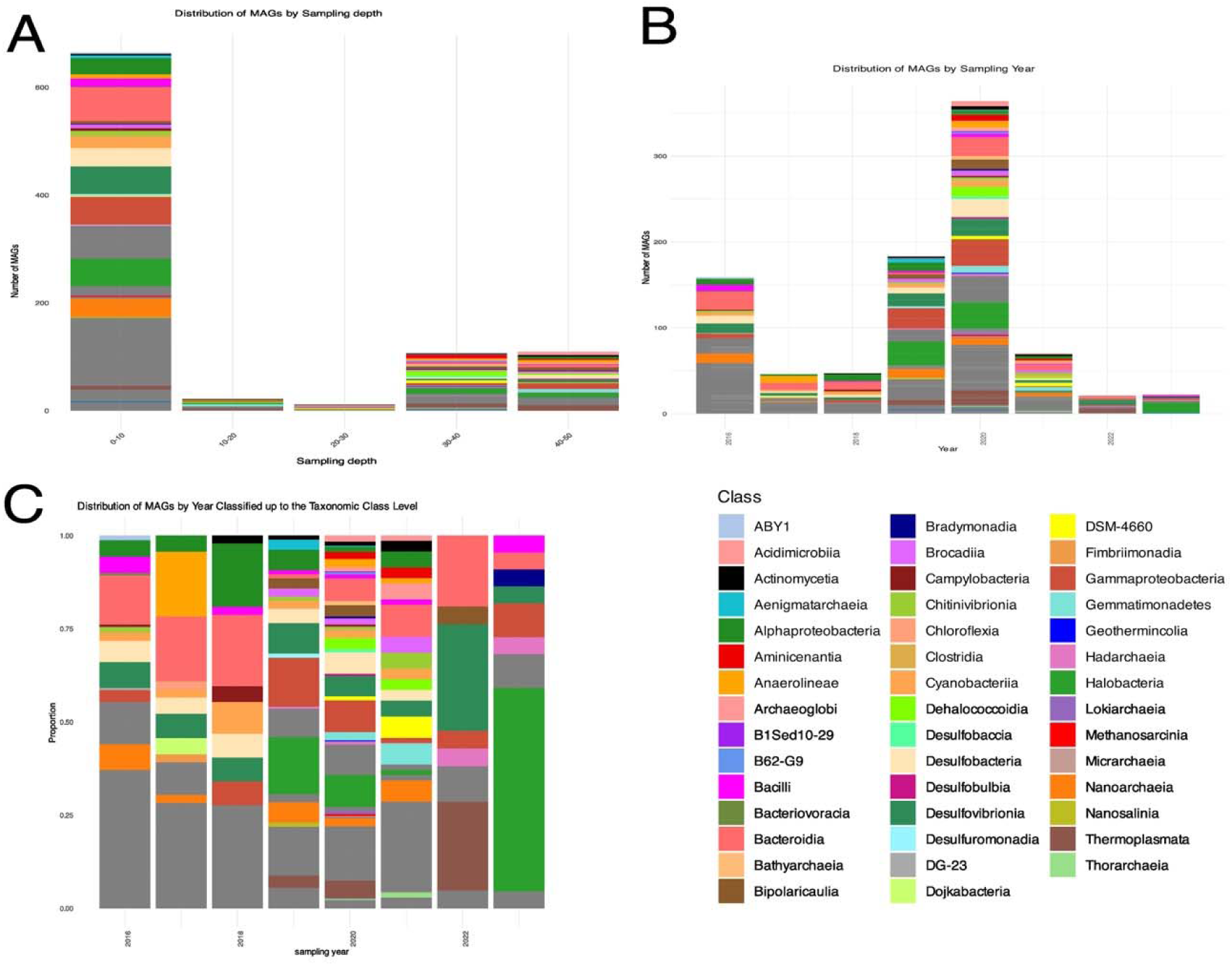
Distribution of MAGs by (A) sampling depth and (B) year. Additionally, the distribution of MAGs by year classified up to the taxonomic class level is shown (C).

**Figure S4.**
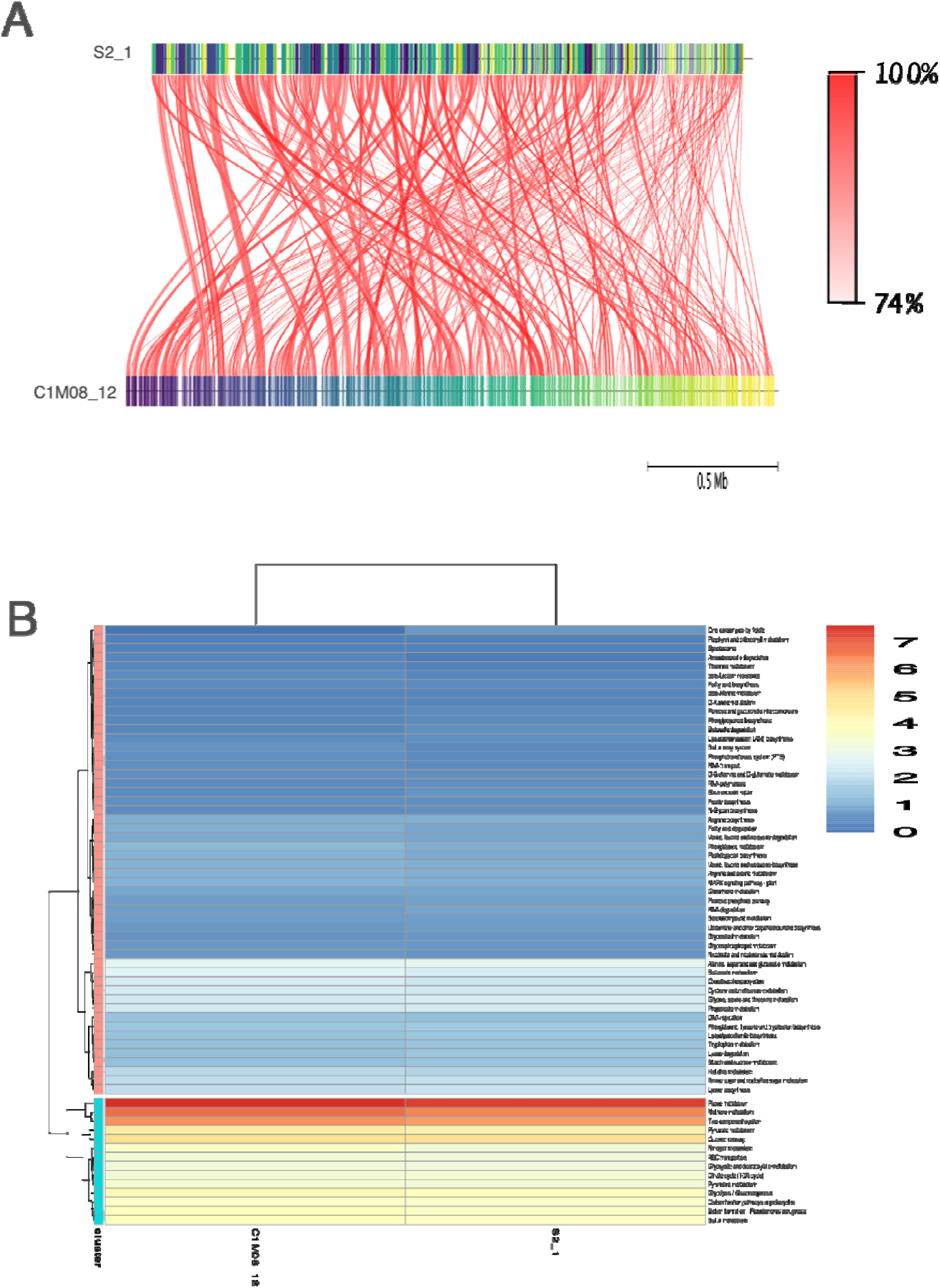
Comparative analysis of MAGs from the genus *Ectothiorhodospira*. (A) Average Nucleotide Identity (ANI) comparison between two MAGs from each site (AD and Churince), focusing on MAGs with the highest completeness and lowest contamination values. (B) Metabolic capabilities of the MAGs, showing no notable differences between them.

